# De Novo TranscriptomeAnalysis Reveals Novel Insights into Secondary Metabolite Biosynthesis in *Tylophora indica* (Burm. f) Merrill

**DOI:** 10.1101/2024.03.05.583525

**Authors:** Anamika Gupta, Deeksha Singh, Rajiv Ranjan

## Abstract

*Tylophora indica* has been widely recognized for its therapeutic properties in traditional Indian medicine. Although its bioactive compounds are used extensively to treat a wide range of ailments, a comprehensive understanding of their genetic basis remains limited. In this study, we conducted a transcriptomic analysis of *T indica* leaf and root using the Illumina platform. High-quality RNA was isolated, and cDNA libraries were constructed for sequencing, generating4.67 GB and5.51 GB of data for leaf and root samples, respectively. 72,795 unigenes and 24,470 coding sequences (CDS) were predicted based on de novo assembly of reads, revealing a complex transcriptome landscape. Functional annotation and pathway analysis revealed biological processes and pathways associated with *T indica*. Based on the Gene Ontology (GO) mapping, the CDS was categorized into biological processes, cellular components, and molecular functions. An analysis of pathways using the KEGG database revealed involvement in critical metabolic pathways. Furthermore, SSRs contributed to the understanding of genetic diversity by identifying simple sequence repeats. In addition, differential gene expression analysis identified genes involved in secondary metabolite synthesis, among other physiological processes. The qRT-PCR validation of selected genes confirmed their differential expression profiles, with roots exhibiting higher expression than leaves. In this study, transcriptomics is conducted for the first time for *T indica*, which may be useful for future molecular research. The detailed findings help us understand *T indica’s* biology, which can be used in biotechnology, and they also show how important it is to protect this species because it is used in medicine.

## Introduction

Ayurveda is a widely used system of traditional medicine in primary healthcare in India. The WHO estimates that 80% of people use herbal treatments worldwide. The demand for medicinal plants has increased as a result of the growing recognition of Ayurveda on a global scale. With 63 known species in the Apocynaceae family, the genus *Tylophora* is extensively dispersed around the world (Sunila and Priya, 2012).

*T indica*, commonly known to as Arkaparni, Indian Ipecac, and Anantmool, is very important in Ayurveda. According to Butani et al. (2007), it has been essential to ethnomedicine and has been used to treat a wide range of illnesses, including cancer, arthritis, bronchitis, diarrhoea, dysentery, cough, and microbiological infections. In particular, *T indica* leaves are used inAyurvedic medicine to treat asthma and bronchitis. *T indica* highlights its medicinal potential due to its abundance of phytochemical elements, including tannins, alkaloids, terpenoids, and saponins (Khare, 2015). *T indica* is a valuable source of bioactive compounds with strong therapeutic potential for treating many illnesses. Many companies have integrated this plant into herbal formulations due to its distinctive qualities, which has piqued the curiosity of researchers in uncovering new medicinal breakthroughs. The widespread usage of *T indica* for therapeutic reasons has caused worries about its preservation status, resulting in its being designated as an endangered plant species in India (Faisal and Anis, 2003; Faisal et al., 2005; Gupta et al., 2020). *T indica* has potential for medical use, but there is still a need for further research on its phytochemical components and the genetic mechanisms involved in their production. The biosynthesis process of significant phytochemicals from *T indica* is not well characterized, especially in terms of the genes responsible for their downstream pathways. Next-Generation Sequencing (NGS) technologies can be used to identify genes related to the biosynthesis pathways of secondary metabolites such as phenolics, alkaloids, terpenoids, and glycosides (Younessi Hamzekhanlu et al., 2022; Tyagi et al., 2022; Singh et al., 2024).

Sequencing the complete genome of medicinal plants is difficult because of the complexity of the genome, high sequencing costs, and computing requirements, leading to a scarcity of fully sequenced genomes. Advancements in transcriptome analysis with Next Generation Sequencing (NGS) have made it easier to understand metabolic pathways in different medicinal plants. Examples include *Artemisia annua* (Paddon and Keasling, 2014), *Wzthania somnifera* (Gupta et al., 2015), *Panax ginseng* (Zhang et al., 2017), *Trichosanthes cucumerina* (Lertphadungkit et al., 2021), *Piper longum* (Dantu et al., 2021), *Psammosilene tunicoides* (Su et al., 2021), and *Tribulus terrestris* (Tyagi et al., 2024). However, there is a lack of transcriptome investigation on *T indica* despite these developments. Deep genome sequencing utilizing Next-Generation Sequencing (NGS) is the primary approach used for generating detailed molecular datasets for identifying proteins in non-model plants such as *T indica*. Exploring *T indica’s* transcriptome extensively using RNA-seq shows great potential in understanding the complexities of this species. The acquired genomic insights have the potential to illuminate the key genes involved in the biosynthesis pathways of important secondary metabolites. Understanding this knowledge improves our understanding of the species’ ecology and evolution and also opens up opportunities for biotechnological interventions (Bohan et al., 2017; Derocles et al., 2018; Kazi et al., 2023).

In this study, the transcriptomes of *T indica* leaves and roots were examined using the Illumina platform. A total of 31,476,234 and 37,163,880 clean reads were obtained from leaves and roots, respectively, resulting in the identification of 72,795 unigenes. Analysis of the transcriptome data revealed gene families associated with secondary metabolite biosynthesis as well as housekeeping genes. This dataset represents the first sequence analysis of *T indica* leaf and root, providing a valuable genomic resource for further molecular investigations.

## Material and Methods

### Collection of plant samples

In September 2018, healthy and disease-free leaf, and root of *Tylophora indica* were collected from the herbal garden of the Dayalbagh Educational Institute located in Dayalbagh, Agra. After each sample was cleaned individually using ultrapure water, it was quickly frozen in liquid nitrogen and kept at - 80°C until RNA extraction.

### Isolation, Qualitative and Quantitative analysis of RNA

Using the PureLink® RNA Mini Kit and following the manufacturer’s instructions, total RNA was isolated from the gathered plant samples. Qualitative analysis of RNA was conducted using a 1% Formaldehyde Denaturing Agarose gel, while quantification was achieved using a Nanodrop 8000 spectrophotometer.

### Illumina cDNA library preparation

Using the instructions provided in the 2xl50 PE Illumina TruSeq Stranded mRNA library preparation kit, lµg of total RNA was used as input for the library preparation process. After enriching mRNA fragments using oligo dT beads, the total RNA was purified, fragmented, and primed for cDNA synthesis. Following fragmentation, mRNA was transformed into first-strand cDNA, which was then amplified using the prescribed number of PCR cycles, A-tailing, adapter-index ligation, and second strand cDNA synthesis. The Agilent DNA High Sensitivity Assay kit was utilized to evaluate the quality of the library.

### De novo Assembly (Mater Assembly)

High-quality reads from Tylophora samples were assembled de novo using Trinity software with the default parameters, which included a k-mer size of 25. Transcripts were represented by fasta sequences that were generated as a result of this approach.

### Unigenes prediction from transcripts

The transcripts underwent additional processing for unigenes prediction using CD-HIT. CD-HIT is a pipeline designed for the analysis of large Expressed Sequence Tags (EST) and mRNA databases. In this process, sequences were initially clustered based on pairwise sequence similarity, and subsequently assembled within individual clusters. This approach aims to generate longer, more comprehensive consensus sequences.

### CDS prediction from unigenes

CDS prediction from the unigenes sequences was conducted using Transdecoder with default parameters. The encoded protein length was set to a minimum of 100 amino acids, and a homology search was performed against the Swiss-Prot and Pfam databases.

### Functional annotation and GO distribution of Predicted CDS/proteins

Using the BLASTX program, the predicted proteins from CDS were compared to the nonredundant (nr) database of the NCBI for similarity searches. GO assignments were utilized in order to illustrate the roles of the anticipated CDS/proteins. Gene product features were represented by ontology-defined terms obtained from GO mapping and classified into three primary domains: Molecular Function (MF), Cellular Component (CC), and Biological Process (BP). For GO mapping, the Blast2GO Pro software was used to get GO terms for all functionally annotated proteins that were acquired by BLASTX against the NR database. A total of 17,015 CDS were assigned at least one GO term, acknowledging that a single CDS could have more than one GO term.

### Pathway analysis of CDS

Ortholog assignment and mapping of the CDS to biological pathways were conducted using the KEGG Automatic Annotation Server (KAAS). All CDS were compared via BLASTX against the KEGG database with a threshold bit-score value of 60 (default). The enriched CDS were categorized into 5 level-I categories and 25 level-2 functional pathway categories. These mapped CDS represented metabolic pathways of major biomolecules such as carbohydrates, lipids, nucleotides, amino acids, glycans, cofactors, vitamins, terpenoids, polyketides, etc. Additionally, the mapped CDS encompassed genes involved in genetic information processing, cellular processes, environmental information processing, and organismal systems.

### SSR (Simple Sequence Repeats) identification

Simple Sequence Repeats (SSRs) or microsatellites are tandem repeats of nucleotide motifs typically ranging in size from 2 to 6 base pairs. These motifs are known for their polymorphic nature and are widely distributed throughout genomes. In this study, SSRs were identified in scaffold sequences using the MISA Perl script. The criteria used for SSR identification were as follows:

- Di-nucleotide pattern should appear at least six times
- Tri-nucleotide pattern >= five times
- Tetra-nucleotide pattern>= five times
- Penta-nucleotide pattern >= five times
- Hexa-nucleotide pattern>= five times

SSRs with flanking regions of 250 base pairs (upstream and downstream) were extracted using an in house Python script, which is also capable of designing primers for SSR amplification.

### Differential Gene Expression Analysis

Differential gene expression analysis was conducted using commonly occurring CDS (based on common NR blast hit accession) for the Leaf-vs-Root combination. To identify differentially expressed genes, reads from leaf and root samples were mapped onto the master assembly CDS sequences using bwa-0.7.5a. The mapped read counts were then used as input for DESeq, an **R** package that provides normalized values in terms of "basemean" for logFC and p-value evaluation. CDS were classified as upregulated if log2FC > 0 and downregulated if log2FC < 0.

For visualization, the top 50 expressed genes, including highly upregulated and highly downregulated genes, were selected. These genes were used to generate a heatmap using the heatmap module of **R** programming.

### QUANTITATIVE GENE EXPRESSION AND VALIDATION

Six genes were chosen at random from the top 50 expressed genes found by heatmap analysis, together with a housekeeping gene, for gene expression confirmation. Using the PureLink® RNA micro kit, RNA was isolated from fresh roots and leaves ofTylophora indica. The extracted RNA was subjected to formaldehyde and MOPS running buffer gel electrophoresis on a 1% agarose gel. The Superscript® III First-Strand Synthesis Kit was used to synthesis cDNA utilizing random hexamers. Primers were created for PCR using the Primer Quest program and diluted to a practical concentration. KAPA SYBR® FAST qPCR master mix (2X) was used for the qRT-PCR. Cycling parameters included an initial denaturation at 95°C for 3 min, followed by 40 cycles of95°C for 30 sec, 55°C for 30 sec, and 72°C for 30 sec. All experiments were performed with three biological and technical replicates. Gene expression levels were normalized using the GAPDH internal control gene and calculated using the 2-^Δ Δct^ method (Livak and Schmittgen, 2001).

## Results

### Quality control and quantification of isolated RNA

The QC evaluation of extracted RNA included analyzing band patterns on a 1% formaldehyde agarose gel for both leaf and root samples, as shown in **Figure 1**. The RNA was quantified using the NanoDrop 8000 spectrophotometer. Both samples exhibit good RNA quality based on the reported specifications.

**Figure 1:**
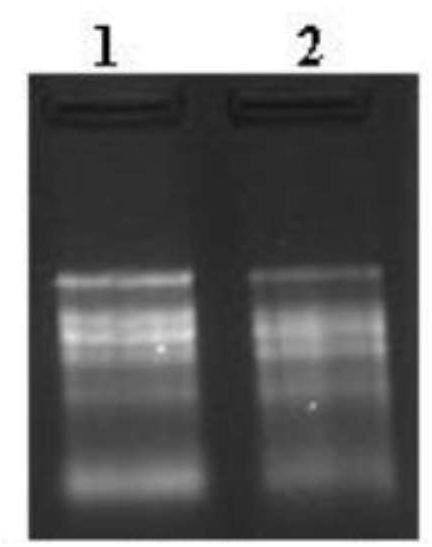
QC of RNA on 1% Formaldehyde Agarose gel. Lane 1: Tylophora fresh leaf and Lane 2: Tylophora fresh root.

After amplification, the library was analyzed using the Bioanalyzer, which showed distinct peaks in the library profiles of both leaf and root samples. This characteristic indicates the production of high-quality libraries. The average library sizes were found to be 399 bp for the leaf sample and 377 bp for the root sample.

### Cluster generation and sequencing

After determining the qubit concentration and mean peak size of the library, it was placed onto the Illumina platform for cluster creation and sequencing. The library preparation process included fragmenting the template and ligating adapters. Paired-end sequencing enabled the sequencing of template fragments in both forward and reverse directions. Library molecules are attached to adapter oligos that are complementary on a paired-end flow cell. Sequencing of leaf and root samples was performed using next-generation sequencing on the Illumina platform using 2xl50 bp sequencing chemicals. 4.67GB of data was generated for the leaf sample and 5.51GB for the root sample. The raw data generated was submitted in the SRA databases of NCBI under the accession number **PRJNA782486**. The leaf sample had a total read count of 31,476,234, whereas the root sample had a total read count of 37,163,880. The leaf sample was sequenced for 4,670,867,015 bases, whereas the root sample was sequenced for 5,517,624,383 bases.

### De novo Assembly (Master Assembly)

De novo assembly is used for sequencing new genomes when no reference sequence is present in databases for alignment. The raw data was filtered using Trimmomatic to remove duplicate sequences. The high-quality readings were assembled de novo using Trinity program with default parameters (kmer 25). The statistical elements of the assembly were computed using in-house Perl scripts, as shown in **Table 1. Graph 1** shows the length distribution of transcripts.

**Table 1:** Statistics of assembled transcripts.

| **Description** | **Master Assembly** |
| --- | --- |
| Total no of transcripts | 87340 |
| Total transcriptome length (bp) | 43772184 |
| Maximum transcript length (bp) | 8554 |
| Transcript N50 (bp) | 694 |

### Prediction ofUnigenes from transcripts

The transcripts were further processed to predict unigenes using CD-HIT. CD-HIT is a specialized tool created for the analysis of extensive EST and mRNA databases. Sequences in this pipeline are first grouped together based on pairwise sequence similarity, and then combined within each cluster to create longer, more comprehensive consensus sequences, which may include quality values. **Table 2** presents a summary of the statistics of unigenes. An analysis was conducted on the distribution of unigenes based on their length. **Graph 2** illustrates the length distribution ofunigenes.

**Table 2:**
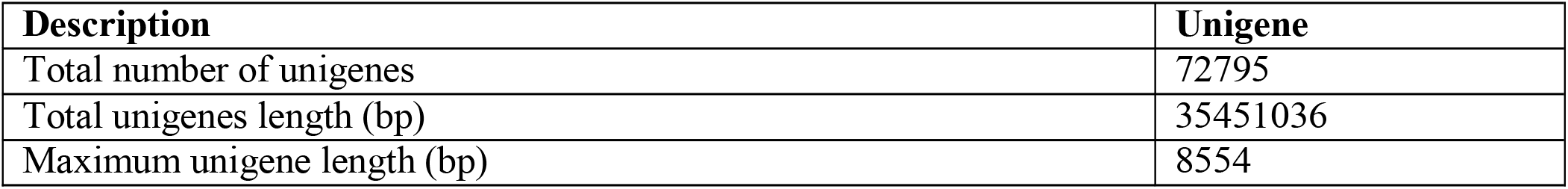
Unigenes Statistics.

### Prediction of CDS from unigenes

Unigenes sequences were used to predict CDS using Transdecoder with default settings. Encoded proteins were required to have a minimum length of 100 amino acids. A homology search was performed using Swiss-Prot and Pfam databases. **Table 3** contains the statistics of CDS. The distribution of CDS was analysed based on their length, as shown in **Graph 3** depicting the CDS length distribution.

**Table 3:** Statistics of predicted CDS.

| **Description** | **#CDS** |
| --- | --- |
| Total number of CDS | 24470 |
| Total CDS length (bp) | 16732368 |
| Mean CDS length (bp) | 683.79 |
| Maximum CDS length (bp) | 4656 |

### Functional Annotation of predicted CDS/proteins

The proteins predicted by CDS were compared to non-redundant NCBI databases using the BLASTX method with an E-value threshold of le-6. **Table 4** displays blast annotation data, whereas **Figure 2** depicts the Venn diagram for functional annotation statistics. The distribution of top-hit species revealed that the majority of hits were found against the plant species *Coffea canephora*, followed by *Sesamum indicum*. **Graph 4** visualizes the distribution of top-hit species after BLAST.

**Table 4:** Blast Statistics of CDS against nr database.

| **Databases** | **Total CDS** | **No. of CDS with Hits** | **No. of CDS with No Hits** |
| --- | --- | --- | --- |
| NR | 24470 | 22828 | 1642 |
| KOG | 24470 | 11573 | 12897 |
| Pfam | 24470 | 10464 | 14006 |
| Uniprot | 24470 | 18533 | 5937 |
| TF | 24470 | 10436 | 14034 |

**Figure 2.**
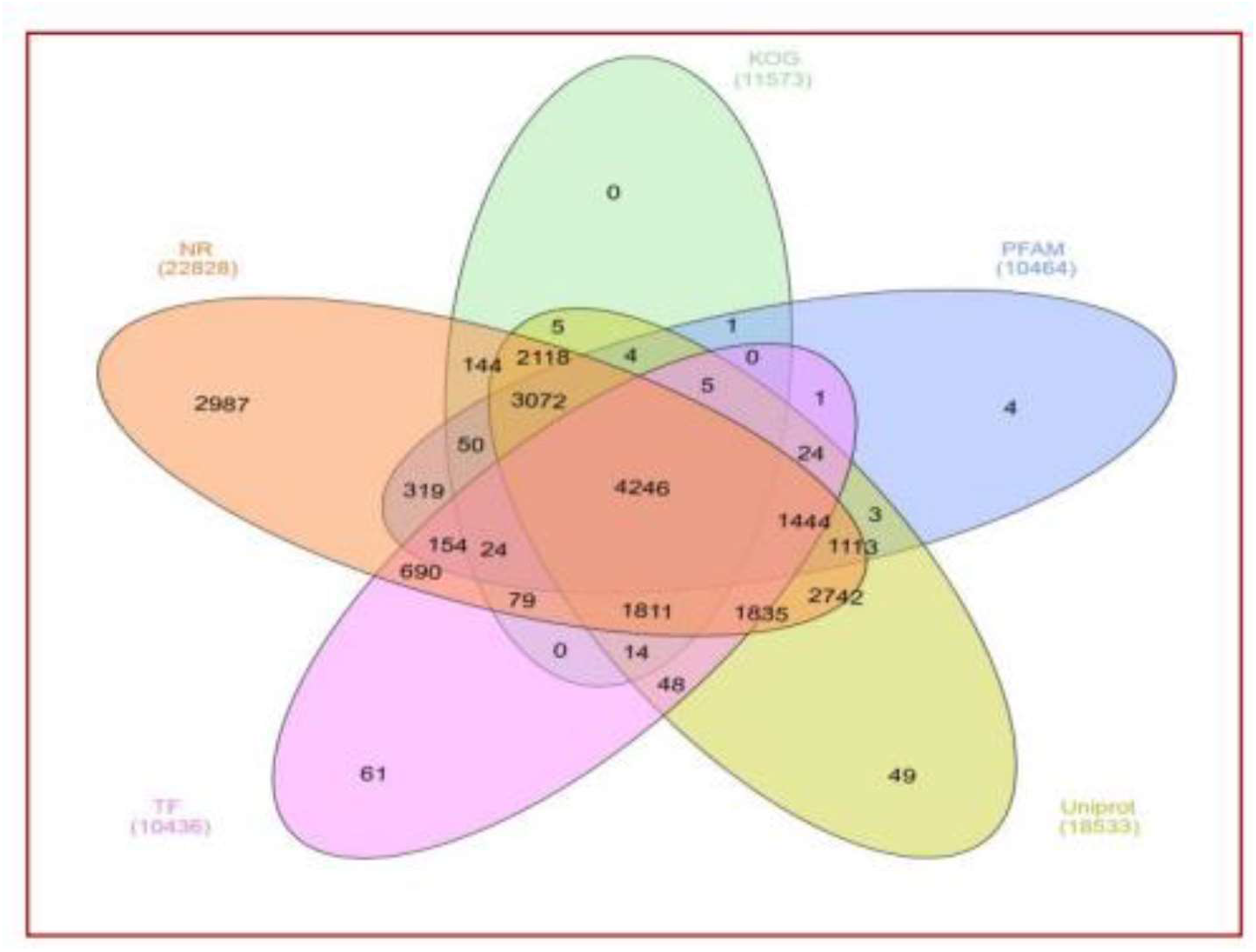
: Venn diagram showing the number of CDS hits in NR, KOG, PFAM, Uniport and TF databases (developed by: Xcleris labs, project id: 999; Date: 31-07-2019)

### GO Sequence Distribution of NR annotated CDS/Protein

The functions of anticipated CDS/proteins were classified using Gene Ontology (GO) assignment, which gives an ontology of defined terms determining gene product characteristics divided into three domains: Biological Process, Molecular Function, and Cellular Components. GO mapping was used to obtain GO terms for all BlastX functionally annotated proteins from the NR database using Blast2GO Pro.

A total of 17,015 CDS were assigned at least one GO term, indicating that a single CDS can have more than one GO term. The distribution of GO categories is as follows: 8,948 CDS were assigned to Biological Process, 6,862 CDS to Cellular Component, and 10,350 CDS to Molecular Function. These CDS were further categorized into sub-categories within these three domains.

In the domain of Biological Processes, a greater number of CDS belong to metabolic processes, followed by cellular processes. In the domain ofMolecular Function, the majority of CDS are associated with binding activity, followed by catalytic activity. Within the Cellular Component domain, the majority of CDS are associated with the membrane, followed by cellular components. **Graph 5** illustrates the distribution of GO terms for all CDS.

### Pathway analysis of CDS

Ortholog assignment and mapping of the CDS to biological pathways were conducted using the KEGG Automatic Annotation Server (KAAS). The CDS were compared with the KEGG database using the BLASTX tool with default parameters, including a threshold bit-score value of 60. The enriched CDS were categorized into 5 level-I categories and 25 level-2 functional pathway categories. These mapped CDS represented metabolic pathways of major biomolecules such as carbohydrates, lipids, nucleotides, amino acids, glycans, cofactors, vitamins, terpenoids, polyketides, etc. The mapped CDS were further divided into various categories, including Metabolism, Genetic Information Processing, Environmental Information Processing, Cellular Processes, and Organismal Systems. Among these categories, 1,261 CDS were involved in signal transduction, followed by 731 in carbohydrate metabolism, 662 in translation, and so on. **Table 5** provides a breakdown of the number of CDS found in each category.

**Table 5:** Category wise distribution of CDS for functional.

| **Metabolism** |  |
| --- | --- |
| Carbohydrate metabolism | 731 |
| Energy metabolism | 364 |
| Lipid metabolism | 393 |
| Metabolism of other amino acids | 168 |
| Amino acid metabolism | 489 |
| Xenobiotics biodegradation and metabolism | 131 |
| Glycan biosynthesis and metabolism | 182 |
| Nucleotide metabolism | 134 |
| Metabolism of cofactors and vitamins | 260 |
| Biosynthesis of other secondary metabolites | 170 |
| Metabolism of terpenoids and polyketides | 139 |
| **Genetic Information Processing** |  |
| Translation | 662 |
| Transcription | 313 |
| Folding, sorting and degradation | 501 |
| Replication and repair | 295 |
| **Environmental Information Processing** |  |
| Membrane transport | 46 |
| Signal transduction | 1261 |
| Signaling molecules and interaction | 1 |
| **Cellular Processes** |  |
| Transport and catabolism | 597 |
| Cell motility | 37 |
| Cell growth and death | 501 |
| Cellular community–eukaryotes | 92 |
| Cellular community–prokaryotes | 60 |
| **Organismal Systems** |  |
| Development and regeneration | 72 |
| Environmental adaptation | 309 |

### Identification of SSR (Simple Sequence Repeats)

The identification of simple sequence repeats (SSRs) in scaffold sequences was carried out using the MISA Perl script. The identification process employed the following parameters: dinucleotide patterns should appear at least 6 times, while trinucleotide, tetranucleotide, pentanucleotide, and hexanucleotide patterns should appear at least 5 times. Descriptive parameters for the identified SSR are presented in **Table 6**. A total of 87,340 sequences were examined, resulting in the identification of 13,100 SSRs. These SSRs were distributed across different unit sizes: 8,356 SSRs were dinucleotide repeats, 4,268 were trinucleotide repeats, 351 were tetranucleotide repeats, 100 were pentanucleotide repeats, and 25 were hexanucleotide repeats. Among these SSRs, 1,225 contained a flanking region of 250 bp, which included both upstream and downstream regions. These SSRs were extracted using an in-house Python script.

**Table 6:** Identified SSR statistics in Master Assembly.

| **Description** | **Master Assembly** |
| --- | --- |
| Total number of sequences examined | 87340 |
| Total size of examined sequences (bp) | 43772184 |
| Total number of identified SSRs | 13100 |
| Number of SSR containing sequences | 10076 |
| Number of sequences containing more than 1 SSR | 2329 |
| Number of SSRs present in compound formation | 1578 |
| SSRs with 250 flanking regions | 1225 |

### Differential Gene Expression Analysis

Differential gene expression analysis was conducted for the combination of leaf-vs-root. Reads from leaf and root samples were mapped onto the master assembly CDS sequences using bwa-0.7.5a to identify expressed genes. In the leaf-vs-root comparison, a total of 9795 downregulated and 9795 upregulated genes were expressed. Among these, 413 were significantly downregulated and 228 were significantly upregulated, based on the log2FC value, where log2FC > 0 indicated upregulation and log2FC < 0 indicated downregulation. Graph 6 illustrates the heatmap for the top 50 expressed genes. In the heatmap, blue colour represents significantly upregulated genes with higher base mean values in roots, while red and yellow colours represent significantly downregulated genes with higher base mean values in leaves compared to roots.

A scatter plot was generated for differentially expressed bile stress genes in leaf-vs-root. Each dot in this plot represents a transcript and its expression value (Graph 7). Dots above the black diagonal line indicate upregulated genes, with red dots representing significantly upregulated genes (p-value < 0.05 and log2FC > 0). Dots below the line represent downregulated genes, with green dots indicating significantly downregulated genes (p-value < 0.05 and log2FC < 0). The volcano plot depicts the distribution of expressed genes based on log2FoldChange and -logl0(pval) **(Graph 8)**. Each dot represents a gene, with red dots indicating a p-value < 0.05; orange dots representing an absolute value of log2FoldChange > 2, and green dots representing a p-value < 0.05 and an absolute value of log2FoldChange > 2. Transcription factors play a crucial role in secondary metabolite biosynthesis. A total of 1073 CDS were grouped with the BHLH (basic helix-loop-helix) family, which was the highest among others. **Graph 9** illustrates the grouping of transcripts with major transcription factor families.

### Validation of selected genes

To validate the gene expression profile, six genes were randomly selected from the top 50 genes of the heatmap, along with one gene chosen as the internal control. Glyceraldehyde 3-phosphate dehydrogenase (GAPDH) was designated as the internal control gene, while the other six genes were Lipoxygenase homology domain-containing I-like, Major Allergen Pro Ar I-like, Chlorophyll a-b binding chloroplastic, RNA-Directed DNA partial, Non-specific lipid-transfer I-like, and NRTl PTR FAMILY. The functions of these selected genes are described in **Table 7**.

**Table 7:**
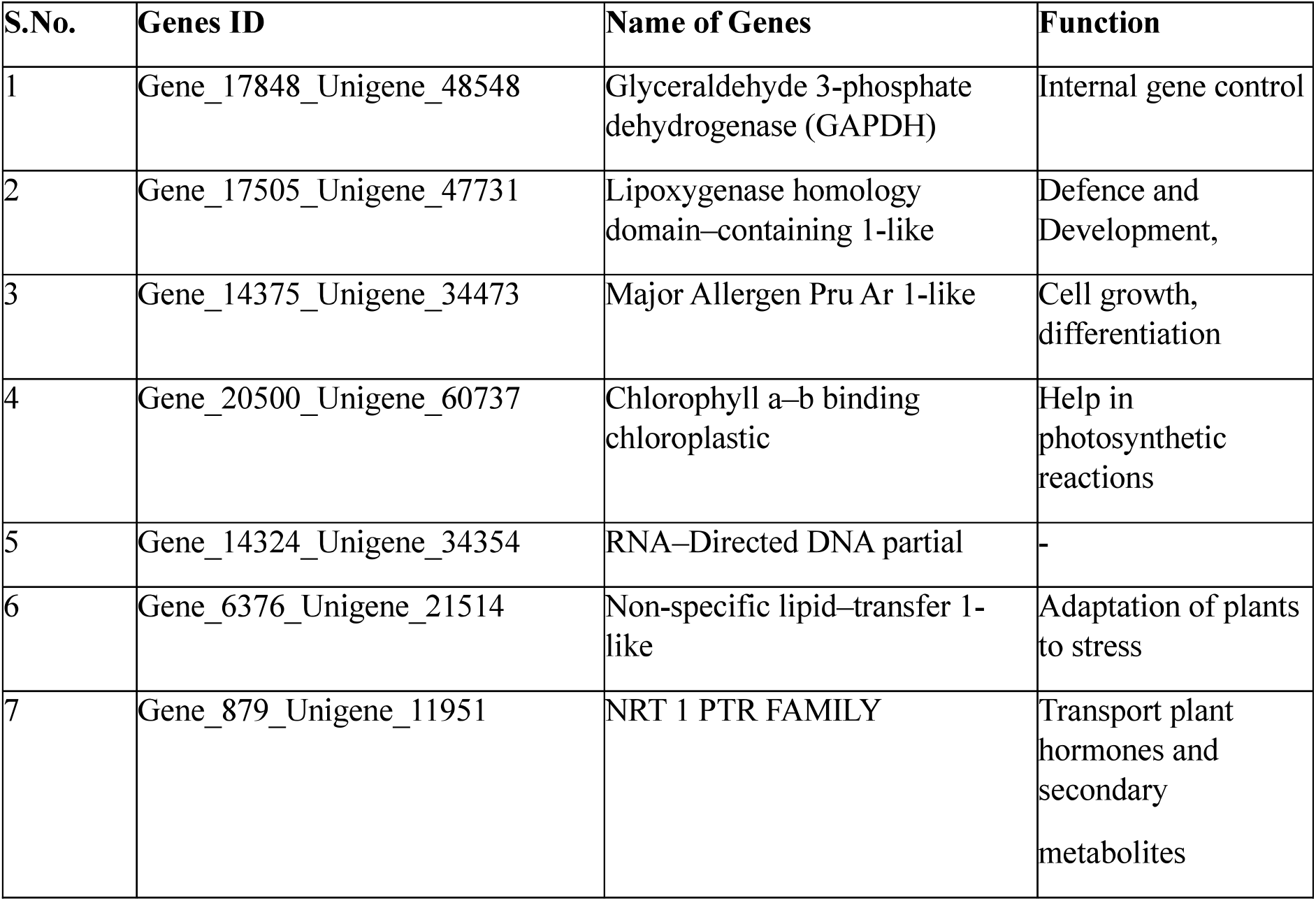
List of selected genes for validation.

Primers for these genes were designed using IDT software, and the list of primers along with the amplicon length is presented in **Table 8**. Leaves and roots of *T indica* were collected for RNA isolation, followed by cDNA synthesis. Expression analysis was performed in terms ofLog2Fold change values, using the GAPDH gene as the internal control to estimate the transcript levels of the genes. All genes were examined with the cycle threshold (Ct) value of the GAPDH gene, and Log2Fold change was calculated. Expression levels were assessed with three biological and three technical replicates. Graph 10 illustrates the gene expression for both upregulated and downregulated levels, with most genes showing higher expression in roots compared to leaves. **Table 9** displays the transcription factor family for the selected genes. These genes were then searched for KEGG pathway analysis, revealing that among the six genes, only two were identified in the KEGG databases. The Chlorophyll a-b binding chloroplastic gene was associated with the KEGG pathway K089 l O LHCA4 (light-harvesting complex I chlorophyll alb binding protein 4), and the gene NRTl PTR family was linked to KEGG pathway Kl4638 SLC15A3-4, PHT (Peptide/histidine transporter).

**Table 8:**
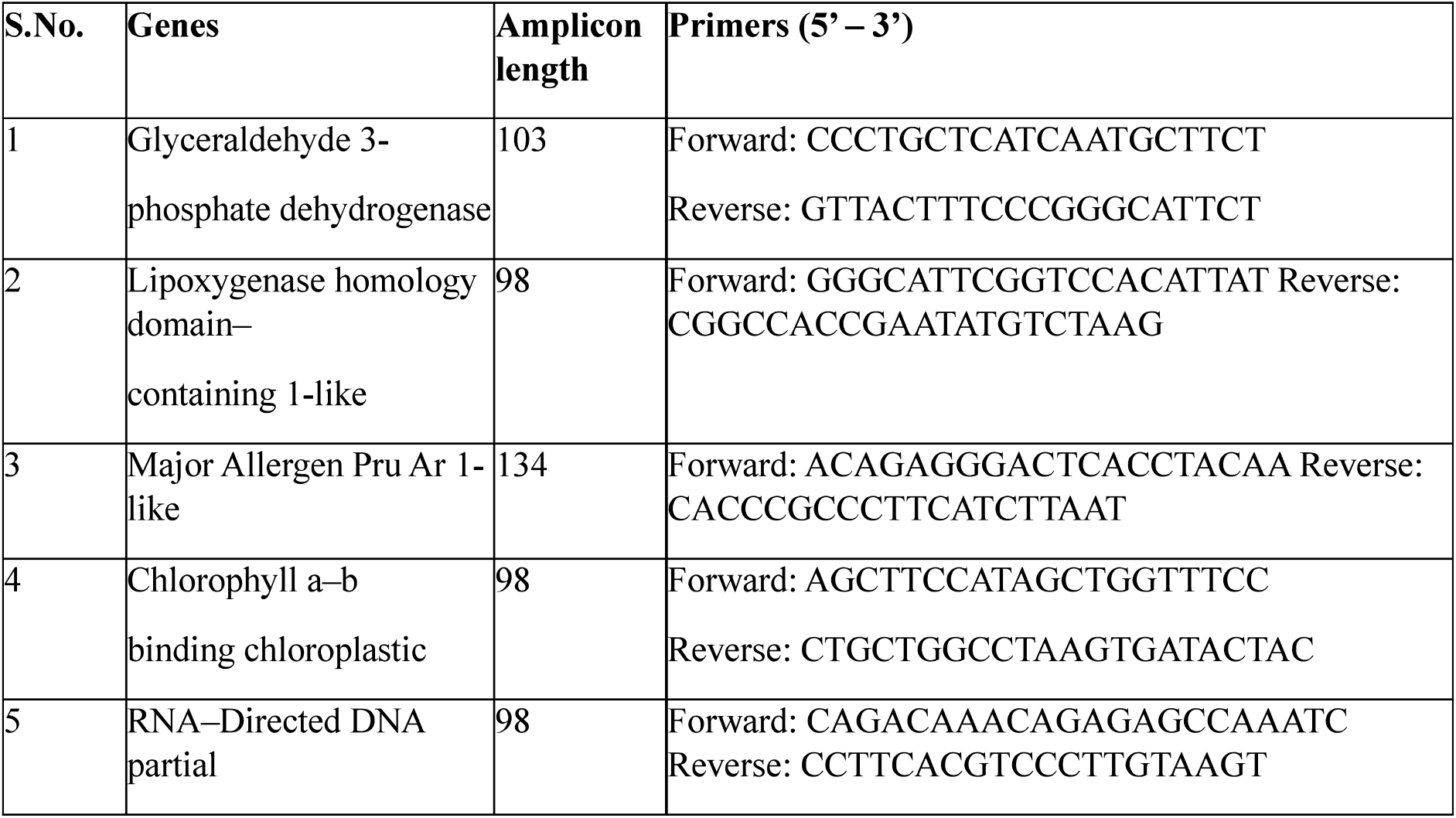
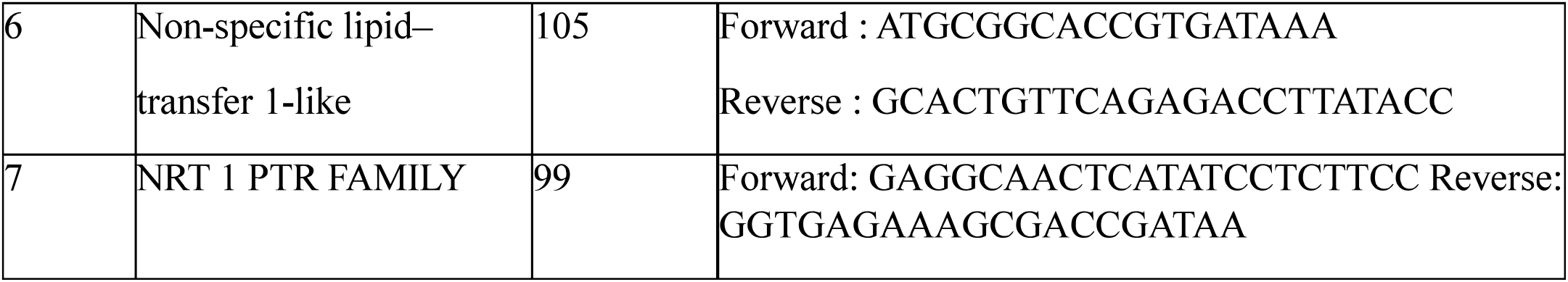
List of primers of selected genes.

**Table 9:**
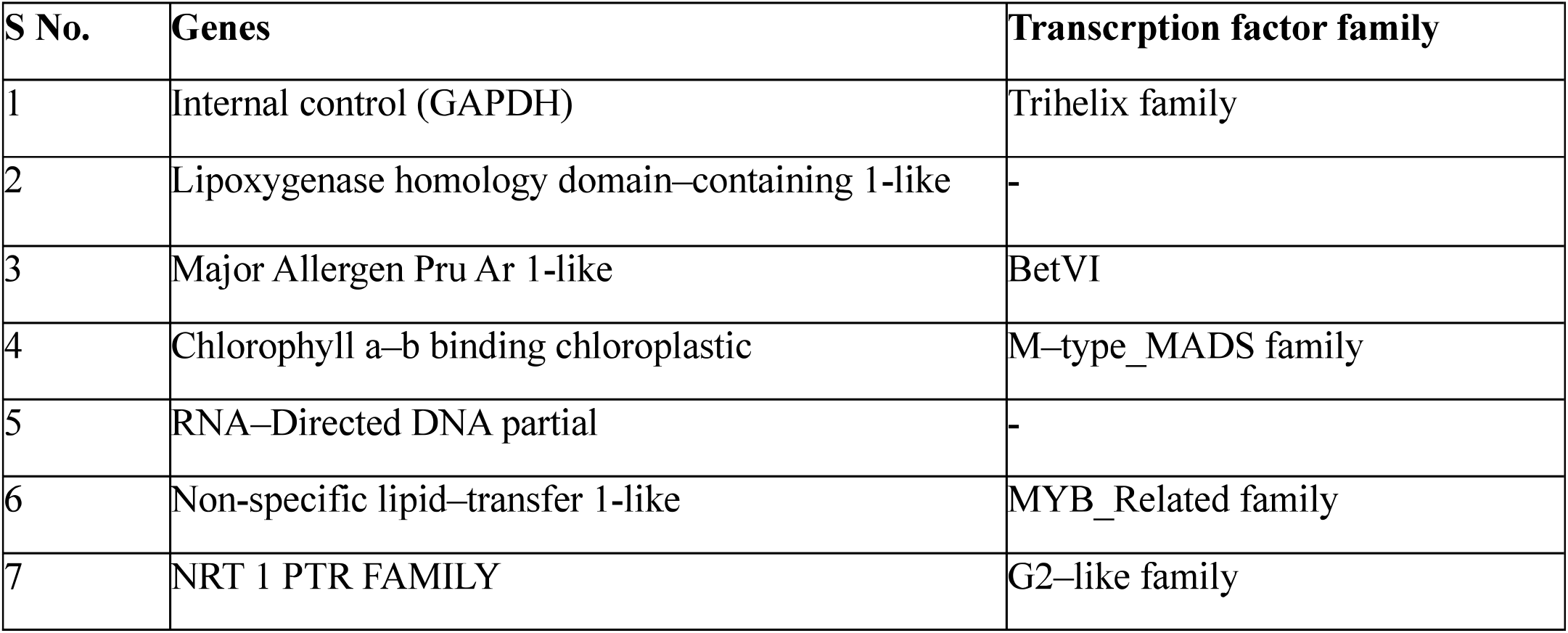
Transcription factor families of selected genes.

## Discussion

The comprehensive analysis conducted in this study provides valuable insights into the molecular characteristics and expression profiles of genes in *T indica*, shedding light on its biological processes and pathways. After RNA isolation, quantification and Quality control was done which shows the good quality of RNA. The initial steps of the study involved the isolation and quantification of RNA from leaf and root samples, ensuring the quality of the extracted RNA through agarose gel electrophoresis and spectrophotometric analysis. Subsequently, the construction of cDNA libraries and high-throughput sequencing using the Illumina platform which generate4.67 GB and5.51 GB data for leaf and root respectively. De novo assembly of reads generate 87340 transcript and 45233 transcript hits on length range >200 & <300, revealing the complexity and diversity of the transcriptome landscape in *T indica*.

Further analyses were performed to predict unigenes and identify coding sequences (CDS) from the transcriptome assembly and 72795 unigenes are predicted. 24470 CDS were predicted from unigenes. Predicted proteins of CDS was search against NR databases using BLASTX and it search against databases NR, KOG, Pfam, Uniport, TF. Highest number of CDS hits with NR databases (22828) followed by Uniport (18533), KOG (l 1573), Pfam (10464) and TF (10436). Top-hit species distribution shows the similarity of *T indica* with *Coffea canephora, Sesamum indicum, Nicotiana tabacum, Nicotiana tomentosiformis, Nicotiana sylvestris, Vztis vinifera, Solanum tuberosum*. GO mapping distributed the CDS into 3 main domains such as biological process with 8948 CDS, Cellular component with 6862 CDS and Molecular function with 10350 CDS. Ortholog assignment and mapping of CDS to the biological pathways were performed using KAAS. Mapped CDS were divided into various categories such as metabolism, genetic information processing, environmental information processing, cellular process, organismal system. Highest CDS hits on signal transduction with 1261 CDS. 731 CDS involved in carbohydrate metabolism and 662 in translation. In soil metagenome, KEGG functional analysis showed that number of orthologs assign to Metabolism category > Environmental information processing > Genetic information processing > Cellular process > Human disease > Organismal systems. In leaf and root transcriptome analysis shows the orthologs assign as Metabolism > Genetic information processing > Environmental information processing > Cellular process > Organismal systems. Differential gene expression analysis was carried out with the combination of leaf-vs-root. It shows expressed gene as 9795 down regulated in which 413 are significantly down regulated and 9795up regulated in which 228 were significantly up regulated on the basis of log2FC value. Transcription factors play a vital role in secondary metabolite synthesis. Major transcription factor families involved in the synthesis, in our study highest number of CDS hits on bHLH family with 1073 CDS. *In Salvia miltiorrhiza* bHLH transcription factor involved in the synthesis oftanshinone biosynthesis (Zhang et al., 2015). For the validation process 6 genes selected from the top 50 significantly expressed genes of heat map. GAPDH was selected as internal control gene or housekeeping gene. Lipoxygenase homology domain-containing I-like, Major Allergen Pru Ar 1 like, Chlorophyll a-b binding chloroplastic, RNA-Directed DNA partial, Non-specific lipid-transfer I like, NRT 1 PTR FAMILY. Lipoxygenase gene family involved in stress response, apoptotic pathway, and signalling (Umate, 2011). Major allergen Pru Ar I like involved in cell growth, Chlorophyll a-b binding chloroplastic helps in photosynthetic reactions, Non-specific lipid-transfer 1-like helps in adaptation for stress in plants, NRT 1 PTR FAMILY involved in plant hormone and secondary metabolites transportation. Expression of genes indicated up regulation as well as down regulation of genes. Among all 6 genes only 2 genes reported in KEGG databases. Chlorophyll a-b binding chloroplastic gene identify for KEGG pathway K08910 LHCA4 (light harvesting complex I, chlorophyll *alb* binding protein 4) and gene NRTl PTR family identify for K14638 SLC15A3-4, PHT (Peptide/ histidine transporter. Transcription factor families belong to gene GAPDH are Trihelix family, Major Allergen Pru Ar 1 like belong to BetVI family, Chlorophyll a-b binding chloroplastic belong to M-type-MADS family, Non-specific lipid-transfer I-like belong to MYB-Related family, NRT 1 PTR FAMILY belong to G2-like family. qRT-PCR studies shows that most of the genes are highly expressed in roots as compare to leaves. Overall, the findings from this study contribute to advancing our understanding of *T indica’s* molecular biology and provide a foundation for further research in plant genomics and breeding.

## Conclusion

The comprehensive analysis carried out in this study provides valuable insights into the molecular characteristics and gene expression profiles of *T indica*, contributing to our understanding of these processes. An analysis oftranscriptomic data from leaves and roots was conducted to identify possible secondary metabolite biosynthesis genes. Genes are highly expressed in the root compared to the leaf based on the differential expression analysis. Among the expressed genes are the Lipoxygenase gene family that plays a role in apoptosis, stress response, and signalling. A major allergen such as Pru Ar 1- is involved in cell growth, a chloroplast gene that binds chlorophyll a-b aids in photosynthesis, a non specific lipid-transfer I-like gene that adapts plants to stress, and a PTR FAMILY that transports hormones and secondary metabolites in plants. Moreover, qRT-PCR validation corroborated transcriptomic findings, suggesting the generated information is reliable and robust. Identifying key genes involved in secondary metabolite biosynthesis and their differential expression patterns provides valuable insight into therapeutic potential and ecological significance of *T indica*. This study contributes to our understanding of *T indica’s* molecular biology and lays the foundation for future research in plant genomics, biotechnology, and breeding. The genomic insights gained pave the way for biotechnological interventions to enhance these valuable plant species’ medicinal properties and conservation efforts.

## Acknowledgments

The authors express their gratitude to the Director ofDayalbagh Educational Institute, Dayalbagh, Agra, for their invaluable support and provision of infrastructure during the research endeavour.

## Figures and Tables

**Graph 1:**
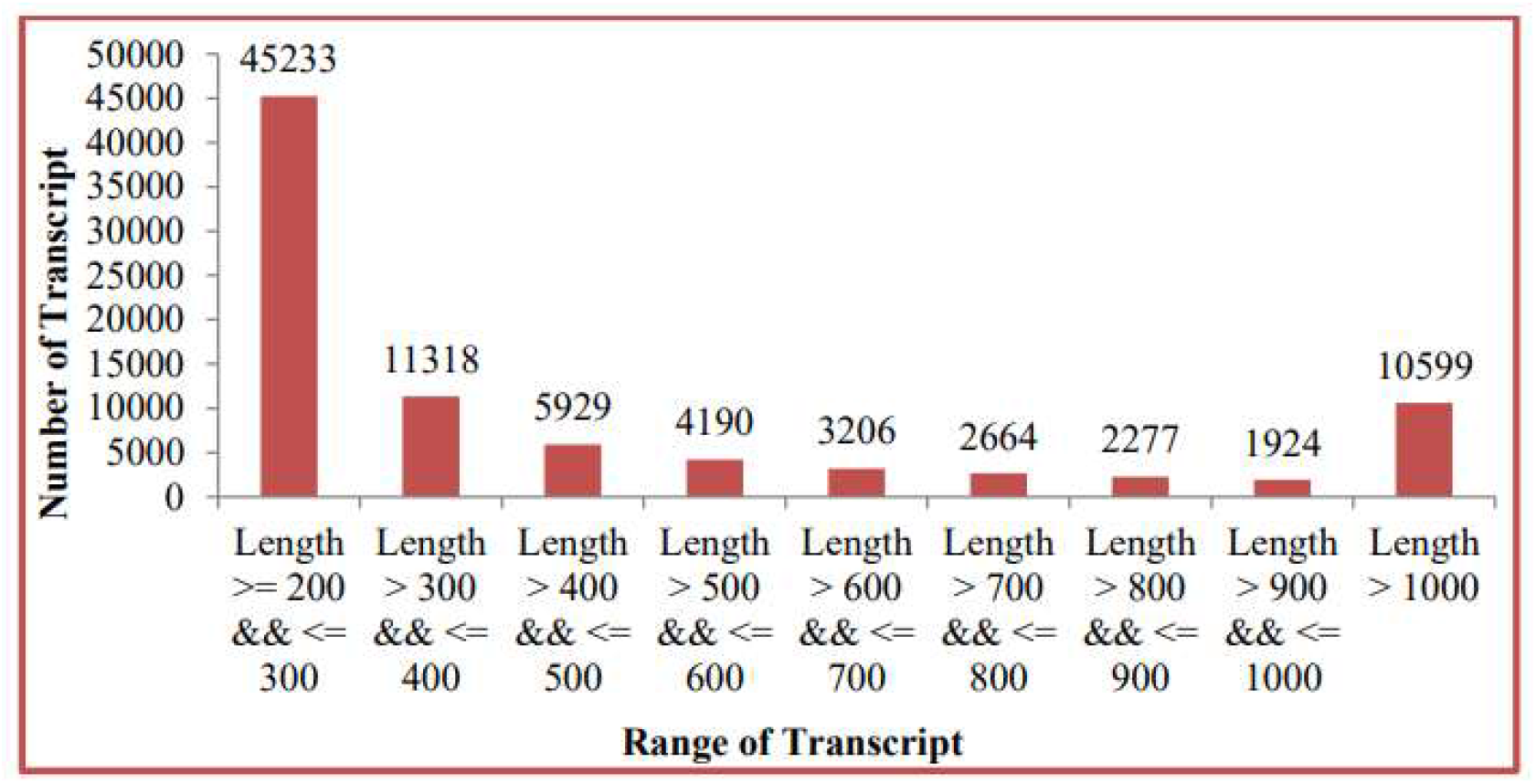
Transcript length distribution where number of transcripts is plotted in Y axis and range of transcript is plotted in X-axis.

**Graph 2:**
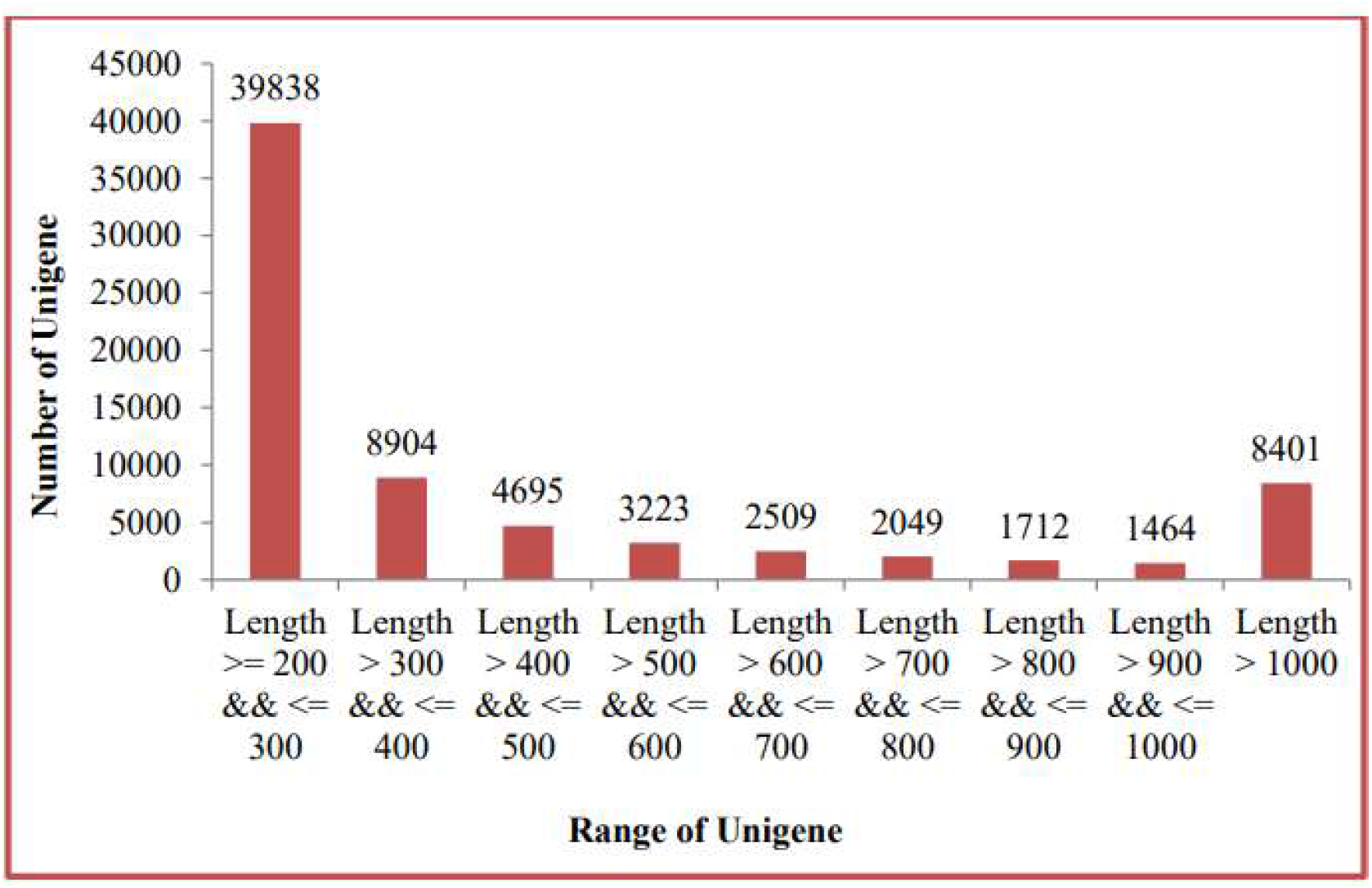
Unigenes length distribution where number of unigenes is plotted in Y axis and range of unigenes is plotted in X-axis.

**Graph 3:**
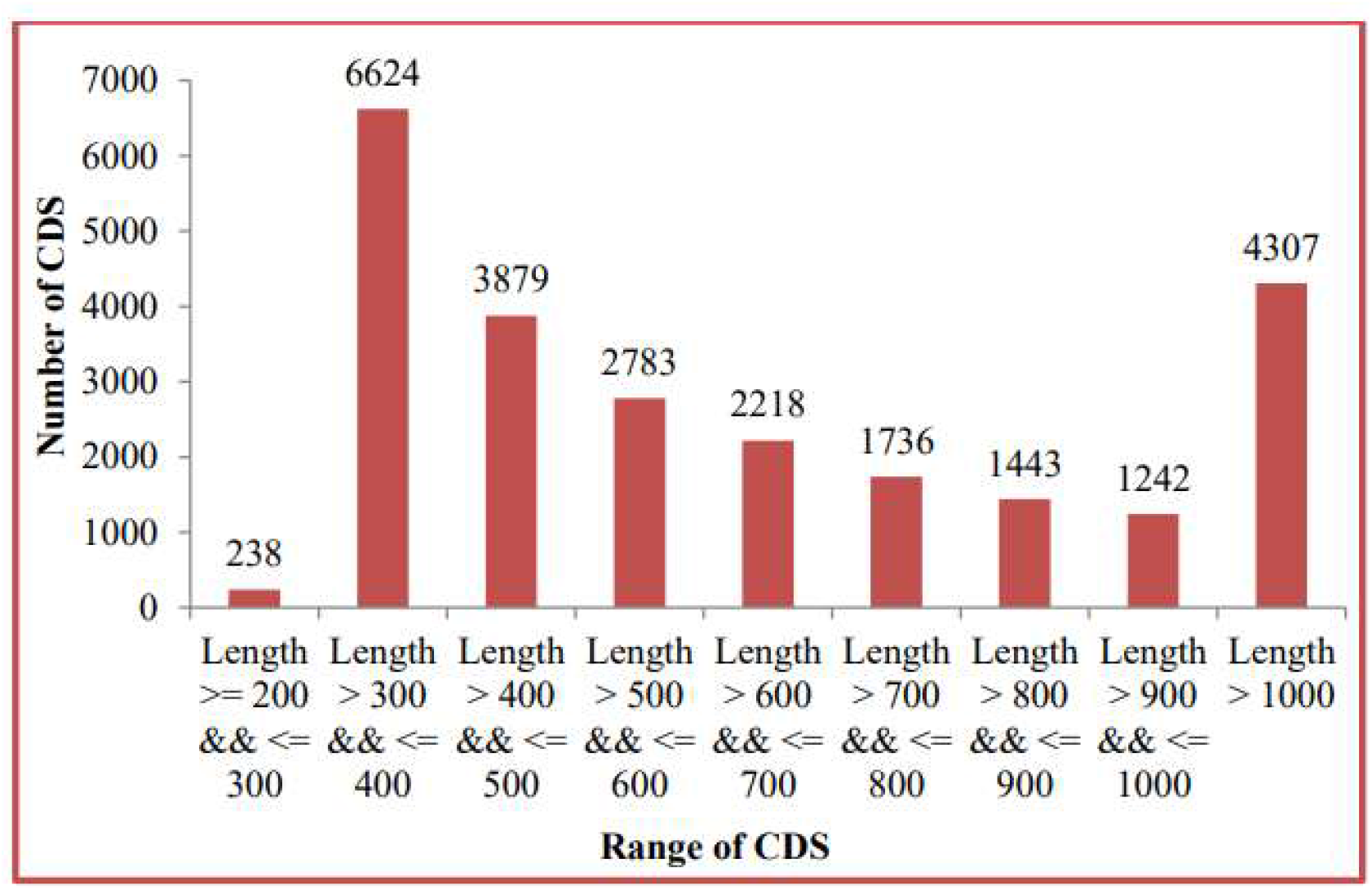
CDS length distribution where number of CDS is plotted in Y-axis and range of CDS is plotted in X-axis.

**Graph 4:**
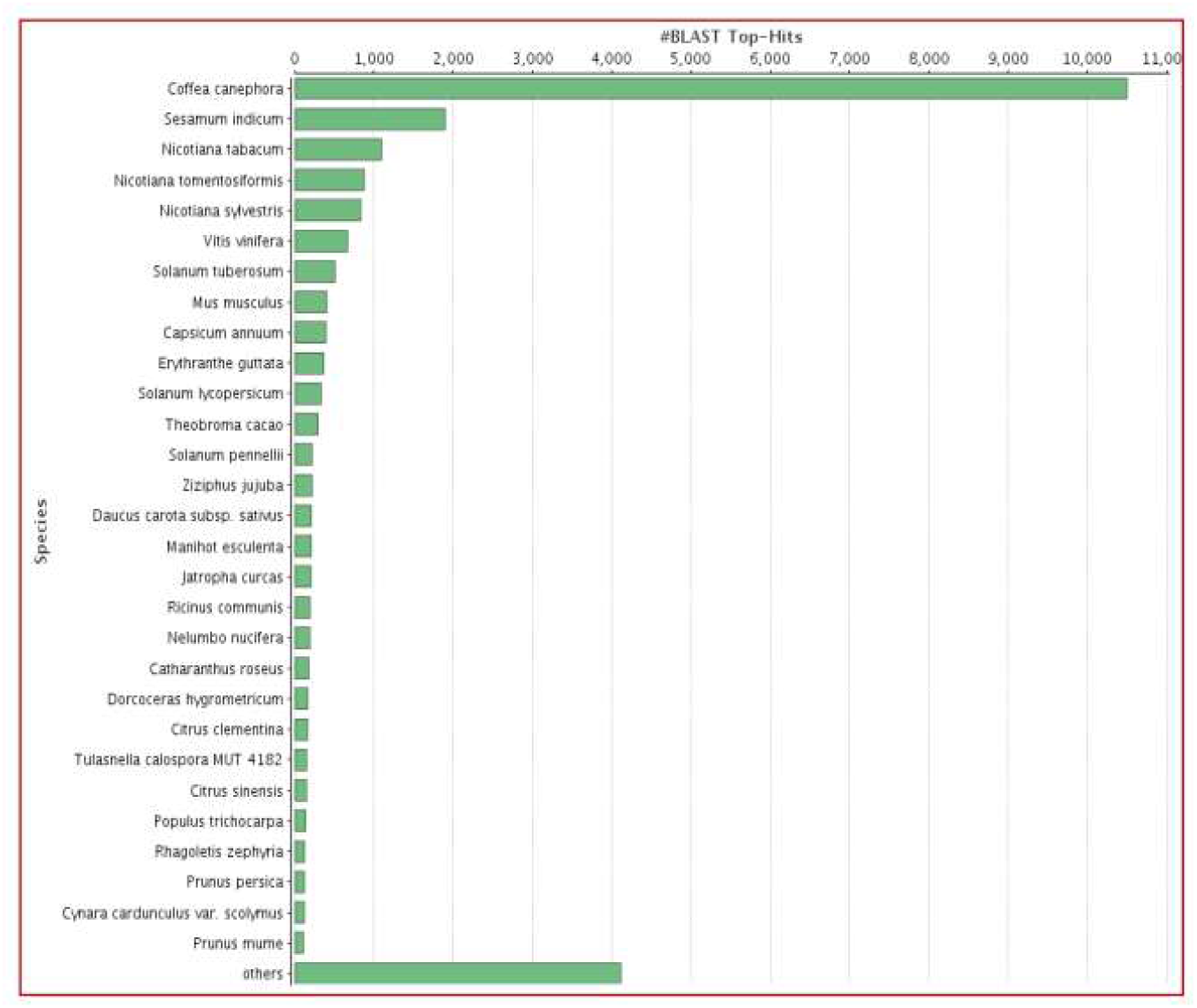
Distribution of CDS in biological process **(BP)**, molecular function **(MF)** and cellular component (CC) (developed by: Xcleris labs, project id: 999; Date: 31-07-2019)

**Graph 5:**
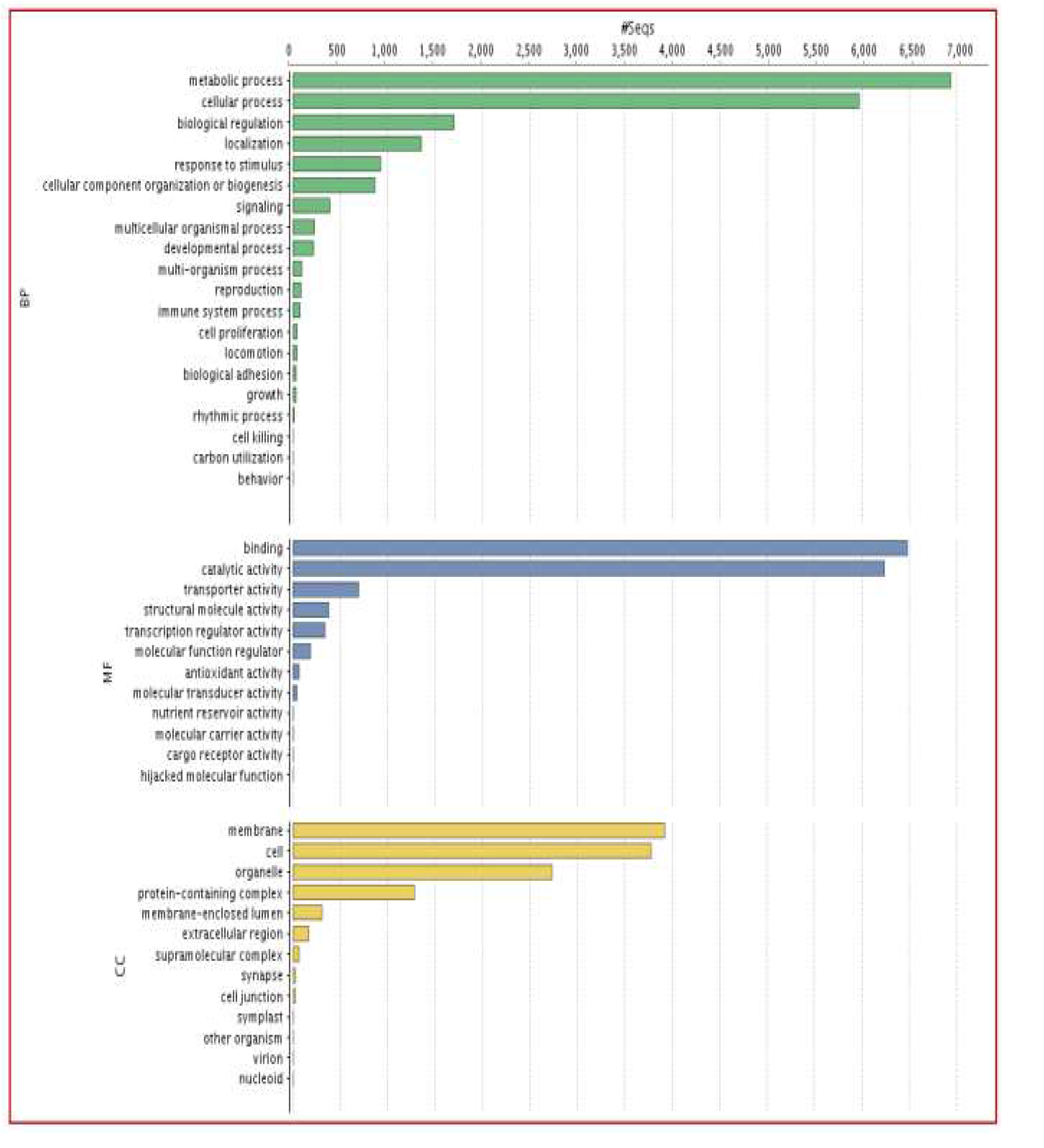
Top-hit species distribution of CDS (developed by: Xcleris labs, project id: 999; Date: 31-07-2019)

**Graph 6:**
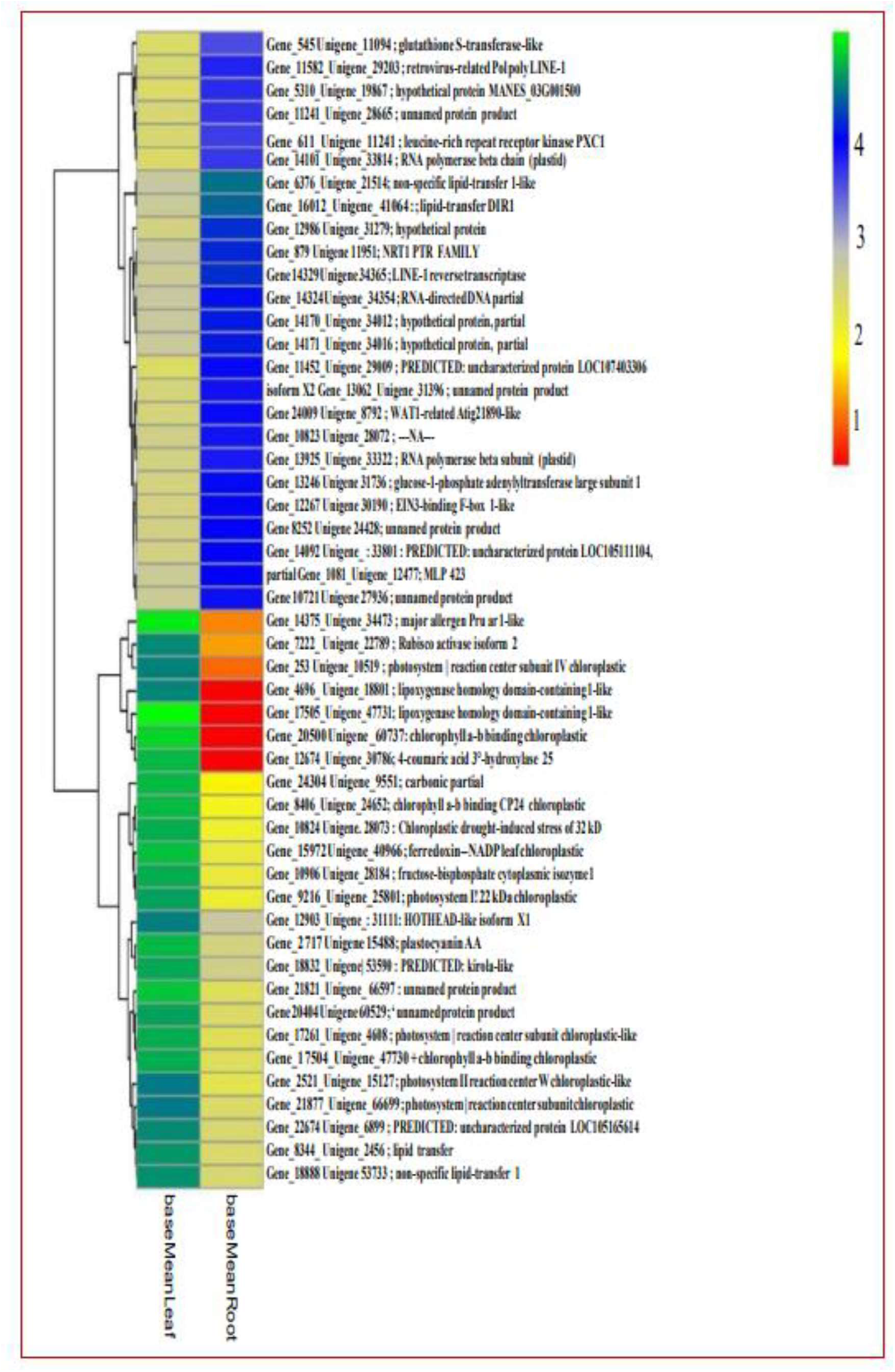
Heatmap analysis for top 50 highly expressed genes.

**Graph 7:**
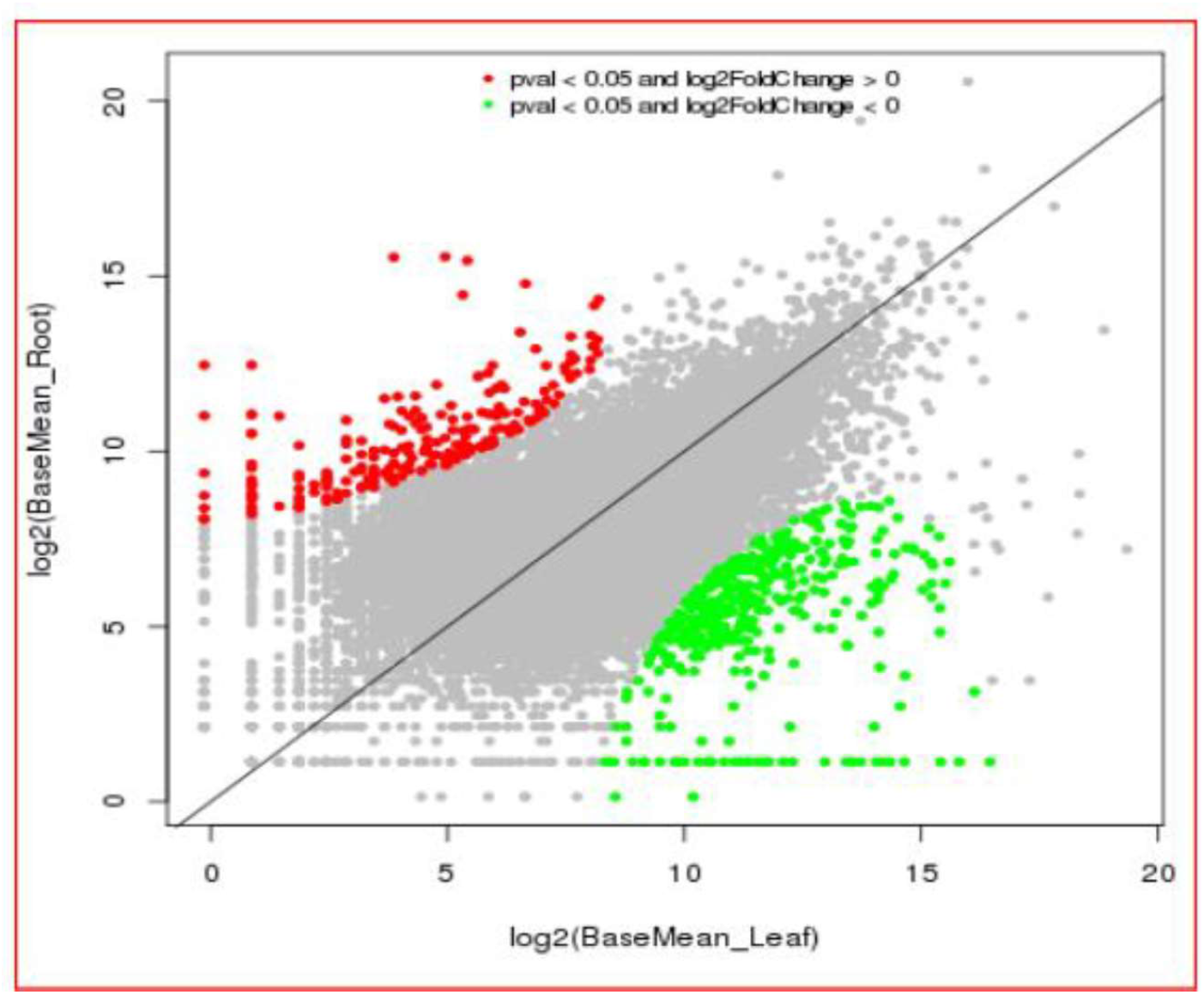
Scatter plot for normalized values obtained through DESeq basemean values of all differentially expressed genes **in** Leaf-vs-Root (developed by: Xcleris labs, project id: 999; Date: 31-07-2019)

**Graph 8:**
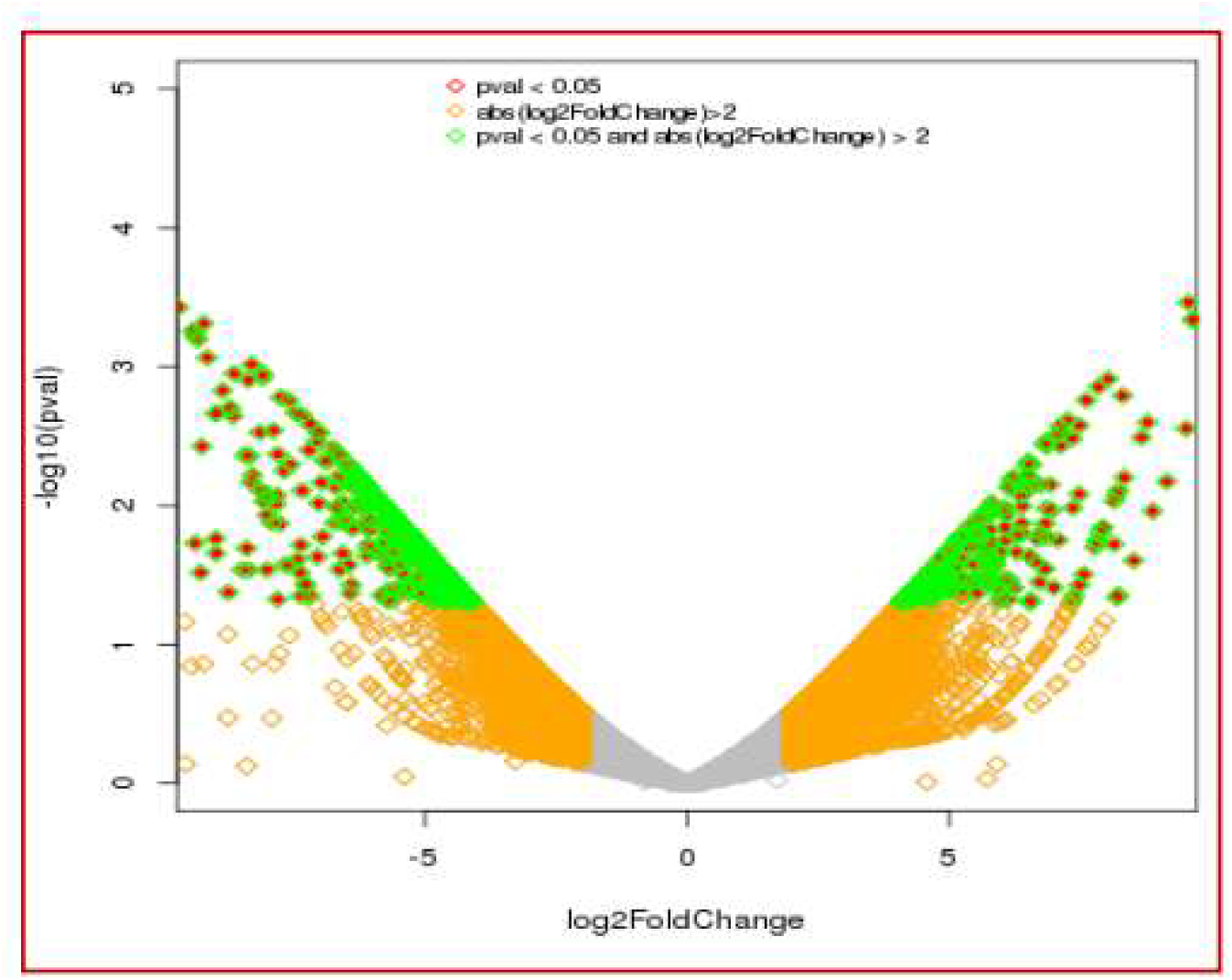
Volcano plots shows distribution of expressed gene for Leaf-vs-Root (developed by: Xcleris labs, project id: 999; Date: 31-07-2019)

**Graph 9:**
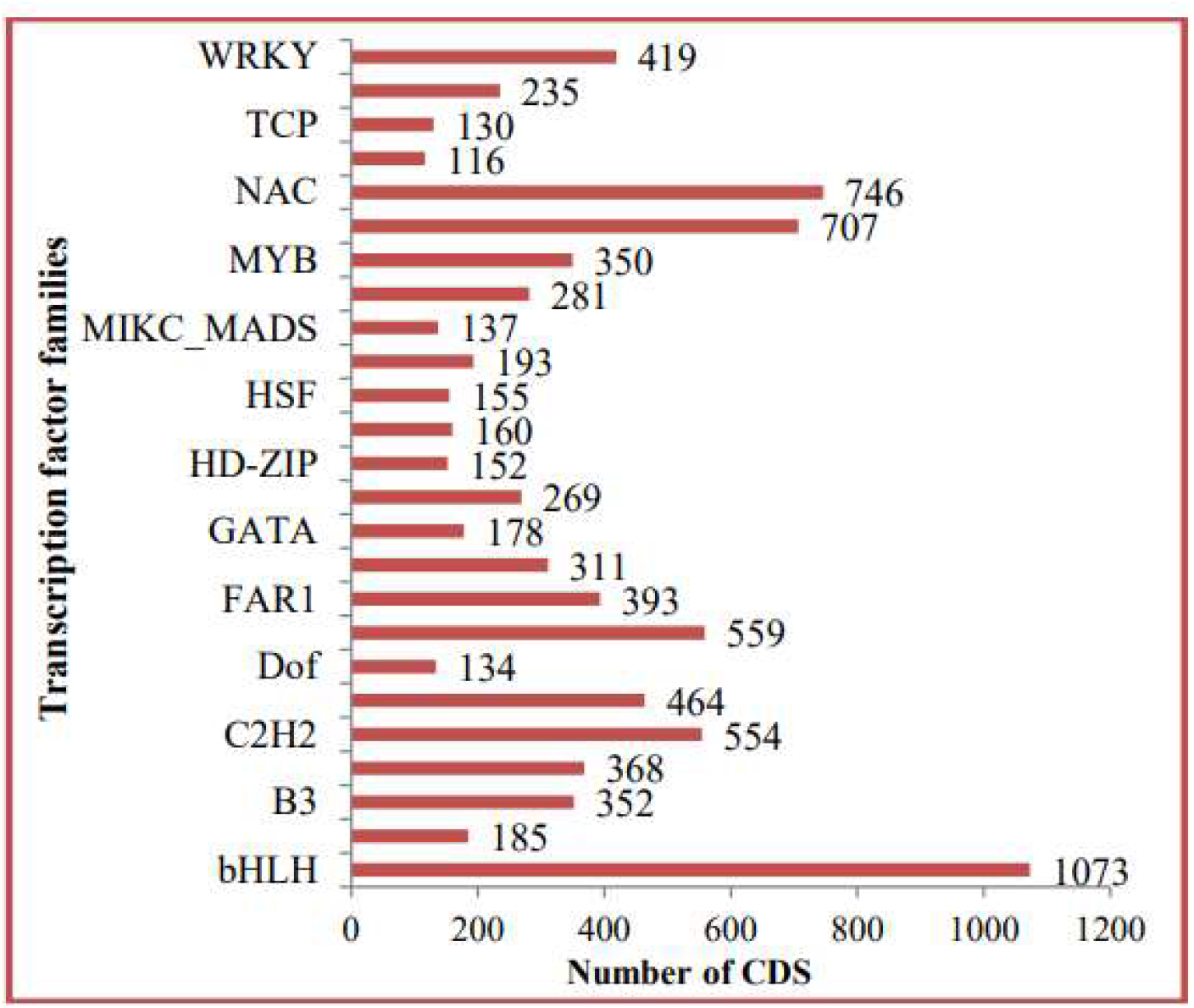
CDS are grouped with major transcription factors families.

**Graph 10:**
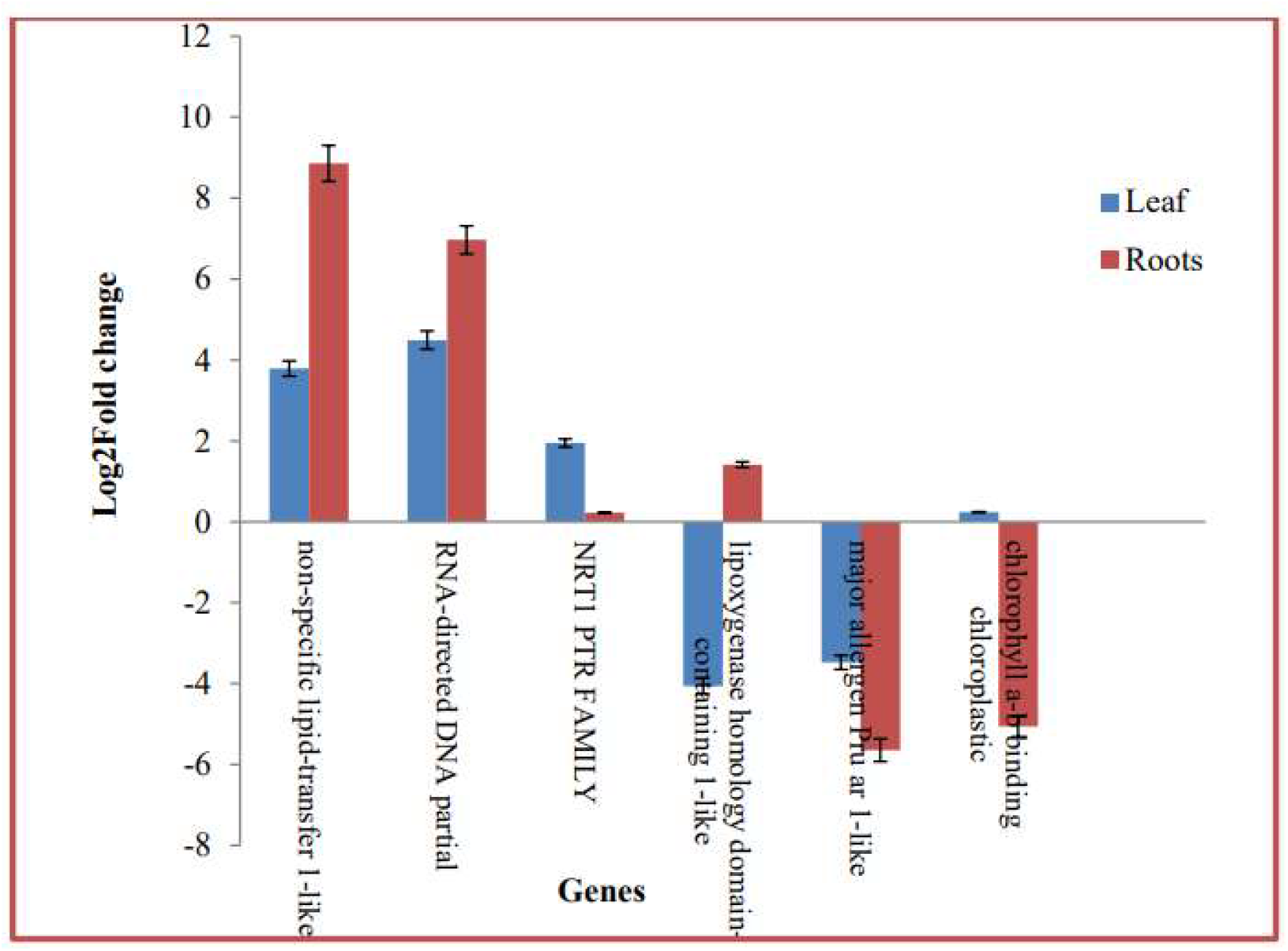
Validation of selected genes through qRT-PCR analysis.

